# Towards Cloud-Native, Machine Learning Based Detection of Crop Disease with Imaging Spectroscopy

**DOI:** 10.1101/2022.12.15.520316

**Authors:** Gloire Rubambiza, Fernando Romero Galvan, Ryan Pavlick, Hakim Weatherspoon, Kaitlin M. Gold

## Abstract

Developing actionable early detection and warning systems for agricultural stakeholders is crucial to reduce the annual $200B USD losses and environmental impacts associated with crop diseases. Agricultural stakeholders primarily rely on labor-intensive, expensive scouting and molecular testing to detect disease. Spectroscopic imagery (SI) can improve plant disease management by offering decision-makers accurate risk maps derived from Machine Learning (ML) models. However, training and deploying ML requires significant computation and storage capabilities. This challenge will become even greater as global scale data from the forthcoming Surface Biology & Geology (SBG) satellite becomes available. This work presents a cloud-hosted architecture to streamline plant disease detection with SI from NASA’s AVIRIS-NG platform, using grapevine leafroll associated virus complex 3 (GLRaV-3) as a model system. Here, we showcase a pipeline for processing SI to produce plant disease detection models and demonstrate that the underlying principles of a cloud-based disease detection system easily accommodate model improvements and shifting data modalities. Our goal is to make the insights derived from SI available to agricultural stakeholders via a platform designed with their needs and values in mind. The key outcome of this work is an innovative, responsive system foundation that can empower agricultural stakeholders to make data-driven plant disease management decisions, while serving as a framework for others pursuing use-inspired application development for agriculture to follow that ensures social impact and reproducibility while preserving stakeholder privacy.

**Key Points:**

- Cloud-based plant disease detection system, easily accommodates newly developed and/or improved models, as well as diverse data modalities.
- Empower agricultural stakeholders to use hyperspectral data for decision support while preserving stakeholder data privacy.
- Outline framework for researchers interested in designing geospatial/remote sensing applications for agricultural stakeholders to follow.

## 1 Introduction

Food security through consistent and scalable agriculture output is the invisible foundation of modern society. However, the globalization and interconnectedness of the agricultural problems we now face demand global, interconnected, and scalable solutions. For example, climate change reduced global farming productivity by 21% over the past 60 years while food demand increased 100% (Ortiz-Bobea et al., 2021). Food demand is anticipated to rise an additional 60-90% over the next 30 years as the population continues to grow (UN, 2015). Plant disease threatens our ability to scale agricultural output to meet this continuously growing demand. Currently, plant disease destroys an estimated 15-30% of the global harvest annually, resulting in $220B in losses (FAO, 2021). Climate change is anticipated to increase the virulence, geographic range, and dispersal capacity of plant pathogens, and consequently, their downstream impacts to human health and society (Nnadi & Carter, 2021). Providing agricultural stakeholders with privacy-preserving, easily deployable, and scalable methods to detect early stage plant disease, the period of time when intervention is both most critical and most likely to succeed, can help address this challenge and reduce food insecurity.

Spectroscopic imagery (SI) in the visible to shortwave infrared light range (VSWIR, 400-2400nm) can quantify chemistry in soil, rock, and vegetation based on the interaction of light with chemical bonds (Curran, 1989). Plant disease changes how solar radiation interacts with leaves, canopy, and plant energy balance, all of which is captured in SI. This underlying capacity is what enables airborne imaging spectroscopy to non-destructively detect biotic stress in both natural and agroecosystems asymptomatically (Zarco-Tejada et al., 2018, 2021; Sapes et al., 2022; Romero Galvan et al., 2022). Disease detection with SI has benefited greatly from data processing using artificial intelligence, specifically machine learning (ML) (Jiménez-Brenes et al., 2019; Hruŝka et al., 2018). However, using ML with SI requires powerful computers and storage media to process exponentially increasing amounts of data. Imaging spectrometers such as the Airborne Visible / Infrared Imaging Spectrometer Next Generation (AVIRIS-NG) instrument have collected SI over millions of acres of agricultural lands unintentionally during campaigns targeted at other uses. Forthcoming satellite systems such as ESA’s Copernicus Hyperspectral Imaging Mission for the Environment (CHIME; (Nieke & Rast, 2018)) and NASA’s Surface Biology and Geology (SBG); (Schneider et al., 2019), will revolutionize global imaging spectroscopy data avail-ability. Taken as a constellation, these instruments will provide data at actionable intervals without cost, and will, for the first time, democratize the availability of such powerful data products for agricultural use. The AVIRIS-NG archives therefore present an exciting opportunity to test and validate the utility of SI for disease management decision making. However, deploying cutting edge models for agricultural stakeholder use beyond academic investigations requires a flexible infrastructure that can readily access an end user’s edge devices, such as computers, storage, and networks. Cloud computing is an ideal solution to this challenge.

Cloud computing has revolutionized data storage and processing by providing ubiquitous, convenient, and on-demand network access to potentially unlimited storage and compute devices (Mell & Grance, 2011). In fact, a third of the world’s data, compute, and storage is hosted by the cloud (Marr, 2015; Cohen, 2021). Practically, the cloud is a collection of servers dispersed in regional or global data centers to provide low-latency computations (e.g. AI and ML) closer to the end users in nearly every industry. For instance, the US National Football League (NFL) relies on Amazon Web Services (AWS) to improve the fan experience and player safety (Amazon Web Services, 2022). In shopping and entertainment, the cloud and ML are the backbone of catalog recommendations or movie recommendations on streaming services. The most profitable segment of the cloud is on-demand access to servers and general-purpose ML models provided by AWS or similar offerings from competitors such as Azure Cloud, Google Cloud Platform, and IBM Cloud (Microsoft, 2022; Google, 2022; IBM, 2022). In agriculture, the cloud and AI are increasingly employed in yield prediction, soil mapping, land management, etc (Analytics, 2022). In the context of SI-informed disease management, cloud computing is not only crucial to scale current methods, but also to explore the implications of future missions. Agricultural decision making stands to benefit from ML models trained using powerful cloud computers. However, in contexts where internet connectivity is sparse, as is often the case in rural communities, it is more practical for models to be trained in the cloud but deployed closer to the data sources. This distributed computing model, which complements cloud computing, is known as edge computing. The edge cloud is a similar compute system organization as the cloud, but rather than providing all computing and storage resources on remote servers, these capabilities are, in part, local and semi-autonomous to accommodate limitations in centralized systems and the intermittency or complete loss of Internet connectivity.

The goal of this work is three-fold. First, we aim to provide a general, cloud-based platform for training and testing SI-informed ML models for non-intrusive and scalable crop disease management. To that end, we showcase new techniques for processing (2.4) and refining spectroscopic imagery (2.3) in a novel agricultural application: grapevine leafroll associated virus complex 3 (GLRaV-3) detection in wine grape (2.1) (Romero Galvan et al., 2022). The second goal is to provide a general platform for researchers to explore the implications of future satellite missions, including SBG and beyond, whose data scales and resolutions are yet to be finalized, for actionable disease detection and decision making. We demonstrate a cloud-based software architecture whose underlying data and application programming interfaces (APIs) are ”plug-and-play” to accommodate current known, and future, yet unknown needs (2.2). The last goal of our work is social impact by making the insights from this work available to agricultural stakeholders, such as grape growers and other industry members, via platform that is only accessible, but also preserves data privacy in mind. To this end, we illustrate the efficacy of ML models developed for GLRaV-3 detection (Figure 6) in a cloud-native environment that does not retain potentially proprietary grower data (e.g. exact field locations) within it. Finally, we discuss methods for reproducibly sharing these models in a manner as required by public funding agencies, while maintaining grower privacy and support (3.1).

## 2 Materials and methods

### 2.1 Model pathosystem, disease incidence, and detection models

Grapevine leafroll associated virus complex 3 (GLRaV-3) is an economically important viral disease of wine, juice, table, and raisin grape. GLRaV-3 is estimated to cause up to three billion (USD) in economic damage to the wine and grape industry annually. While the virus is primarily vectored by mealybugs (Pseudococcidae sp.), it can also be spread by many other phloem feeding insects (J. G. Charles et al., 2009; J. Charles et al., 2006; Golino et al., 2002; Pietersen et al., 2013). Vines infected with GLRaV-3 ultimately have reduced lifespan and productivity. Additionally, infection causes uneven berry ripening and disordered berry chemistry, which reduces wine quality. Detecting GLRaV-3 infected grapevine at the symptomatic stage is straight forward: scouts are trained to recognize the foliar symptoms 1 and a piece of the grapevine is sent over to a commercial lab to verify viral presence via serological or molecular testing. However, the main challenge growers face is early detection. Infected plants can stay at the asymptomatic stage for up to one year, meaning they display no visible sign of disease for humans to identify despite being infectious to nearby grapevines (Maree et al., 2013; Almeida et al., 2013; Naidu et al., 2014). Additionally, unlike red grape varieties (e.g. Cabernet Sauvingnon), white grape varieties (e.g. Chardonnay) do not manifest visible, foliar symptoms that can be identified by scouts (Naidu et al., 2014; J. Charles et al., 2006; Olmos et al., 2016). Existing methods for asymptomatic GLRaV-3 rely on molecular testing which can cost between $40-$300 USD per vine depending on the number of viruses that are being tested. Considering that a small vineyard has close to a 1,000 vines, and a large vineyard 30,000+, this presents a crucial scaling problem imaging spectroscopy is ripe to address.

All data worked with here is within the city of Lodi, California, USA. Here, there are roughly 11,000 acres of vineyards where AVIRIS-NG acquisitions were made. The AVIRIS-NG acquistions captured vineyards roughly a week after the disease incidence coordinates were recorded. For more detailed description of disease incidence collection, validation, and aggregation, see Romero Galvan (Romero Galvan et al., 2022). In brief, disease incidence data was collected at seven vineyards and covered 7 acres (ac) of red grape variety Aglianico, 204ac of Cabernet Sauvignon, and 57ac of Petite Sirah in the city of Lodi, CA by visual inspection for foliar symptoms (”scouting”) by expert teams with a subset sent for external validation via molecular testing in September of 2020 and 2021. Molecular testing found the expert scouts to be 100% accurate. When a diseased vine was encountered, scouts recorded the coordinates of disease incidence in local Universal Transverse Mercator geographic coordinate system, Zone 11 (EPSG: 26910). Each disease incidence point is classified according to one of three labels: non-infected (NI), symptomatic (Sy), and asymptomatic (aSy). It is well established that GLRaV-3-infected red variety grapevines (e.g. Cabernet Sauvignon, Malbec) can stay at the asymptomatic stage for up to a year if no visible symptoms appear before a grapevine’s winter dormant period (J. Charles et al., 2006; Olmos et al., 2016). Therefore, incidence points identified as visibly infected in 2021 were labeled asymptomatic (aSy), and vines identified as visibly diseased by the scouts in 2020, during the AVIRIS data acquisition, are labeled as symptomatic (Sy). Vines that were identified as Sy in 2020 were removed prior to the 2021 growing season, and therefore were not present at the time of 2021 scouting. Vines that were not identified as visibly infected in either 2020 or 2021 are labeled non-infected (NI). We note that we did not conduct molecular testing to prove that these vines were truly asymptomatically-infected at the time of data collection, as it would have been unfeasible given our scope of study and sample testing expense ($40-50 per vine). We emphasize that this assumption is well-supported by current understanding of disease biology (Maree et al., 2013; Almeida et al., 2013; Naidu et al., 2014), and the fact that all green foliage was destroyed and removed from the vineyard during mechanical harvest soon after the flight took place. This implies a lower likelihood that vines experienced an opportunity to become infected between the time of the AVIRIS-NG flight and bud break the following season.

Following common convention of training ML models (Gholamy et al., 2018), our Random Forest models (RFm) were trained through a 70/30 training/validation split. The models were trained on spatially-resampled three-meter SI as Romero Galvan found to be optimal for GLRaV-3 detection in grapevine(Romero Galvan et al., 2022). In total, 268 acres were classified capturing 80% of disease incidence in the field.

### 2.2 Software Architecture

The software architecture is comprised of three major components that provide an adaptable pipeline for disease detection. These components are the NASA Cloud, the experimental lab at our university, and the publicly accessible cloud, specifically, Azure Cloud (Figure 4). The NASA Cloud serves as the peripheral location of raw data. The raw data is collected through NASA’s Airborne Visible/Infrared Imaging Spectrometer Next Generation (AVIRIS-NG) flight missions. That is, the servers host high-dimensional, spectroscopic image (SI) data originating from flights over 875,000 acres of vineyards in California, USA. The SI is accessible via a user-initiated download of Excel spreadsheets containing download links to each image. These datasets serve two distinct functions. First, they serve as data to train new disease detection models. Secondly, they function as raw data for filtering during field-specific inference requests. The experimental lab serves as a central location for data pre-processing and initial model training and testing. Before the NASA Cloud data can yield useful insights, it requires extensive pre-processing to account for noise in the data collection process, as illustrated in Figure 5. In addition to data pre-processing, the experimental lab serves as training ground for the GLRaV-3 model training and testing. The local training and testing follows from three practical considerations. First, it affords researchers as much flexibility as possible in setting up their environments in terms of software dependencies. Secondly, to minimize cloud consumption costs, disease inference requests can be iteratively tested locally before being deployed to the cloud. Lastly, though rare in research labs, the connectivity to the public cloud for data and model access may be limited, and the local setup may intermittently serve inference requests during such outages.

### 2.3 Spectroscopic Imagery

Spectroscopic Imagery (SI) from NASA-JPL’s AVIRIS-NG was used to train these models. Specifically the SI image IDs were: ang20200918t210249, ang20200918t205737, ang20200918t212656, ang20200918t213801, ang20200918t213229. The SI imagery was collected on September 18th, 2020 between 1:00 PM and 3:00 PM Pacific (local) time. All listed SI are collected at the one-meter spatial resolution, collected spectral-channels at 5-nanometers (nm) between 380nm and 25000nm totaling 425 channels (Chapman et al., 2019; Thompson et al., 2018). All AVIRIS-NG imagery used in this study is publicly available reflectance data that can be downloaded from the AVIRIS-NG data portal (https://aviris.jpl.nasa.gov/dataportal/). All AVIRIS-NG SI is atmospherically corrected using the ATmospheric REMoval (ATREM) algorithm (Gao & Goetz, 1990). Additionally, water absorption feature wavelengths are excluded due to noise, specifically, the spectroscopic ranges excluded are: 380nm - 400nm, 1310nm - 1470nm, 1750nm - 2000nm, and 2400nm - 2600nm. All pixels outside the extent of the vineyard are likewise excluded by masking the imagery using the python package Rasterio (Gillies et al., 2013–). To exclude non-vine spectra from the imagery (e.g. soil, shadows, trees), a vine mask was generated using spectral unmixing residuals methodology (Sousa et al., 2022). Both bidirectional reflectance distribution function (BRDF) and topographic correction (Queally et al., 2022) scripts from the HyTools package (Chlus et al., 2022) were applied to all SI. A subset of the AVIRIS-NG imagery required further spatial corrections to improve spatial alignment with the disease incidence scouting data. The National Agriculture Imagery Program (NAIP; USDA) imagery collected within a week of our AVIRIS-NG imagery was used as a reference to improve georeferencing and co-registration. Ground control points (GCPs) were generated by passing a target SI and reference NAIP image to the open-source python library Automated and Robust Open-Source Image Co-registration Software (AROSICS) (Scheffler et al., 2017).

### 2.4 Processing SI Data

Our pipeline takes AVIRIS-NG Imagery with the corrections in Figure **5B** already applied. The automatic pipeline is one that takes AVIRIS-NG Imagery and outputs a classified geospatial raster. The pipeline steps are illustrated in Figure **3A-F**. The first step masks the AVIRIS-NG SI to the extent of the boundaries supplied by an end-user. Spectral residual weights are then used to generate a vegetation mask **3C-D**, the resulting mask is used to remove pixels where the spectral signal of vegetation is at least 50% of the total spectral contribution (**3F**). The resulting masked image is then classified using the RFm generated in section 2.1.

**Figure 1.**
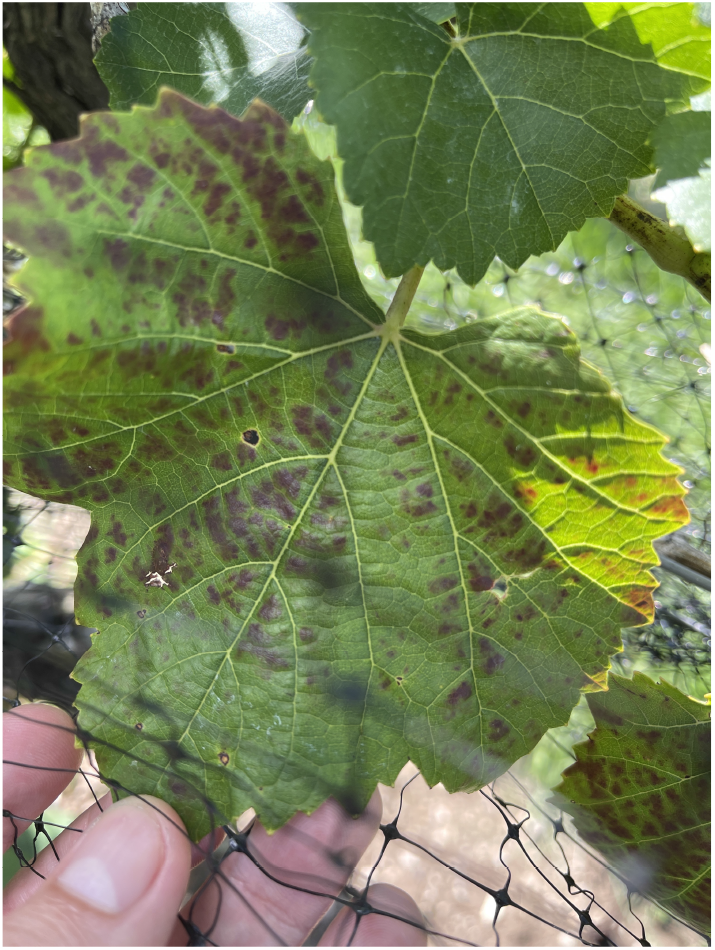
GLRaV-3 infected grapevine of Cabernet Sauvignon variety.

**Figure 2.**
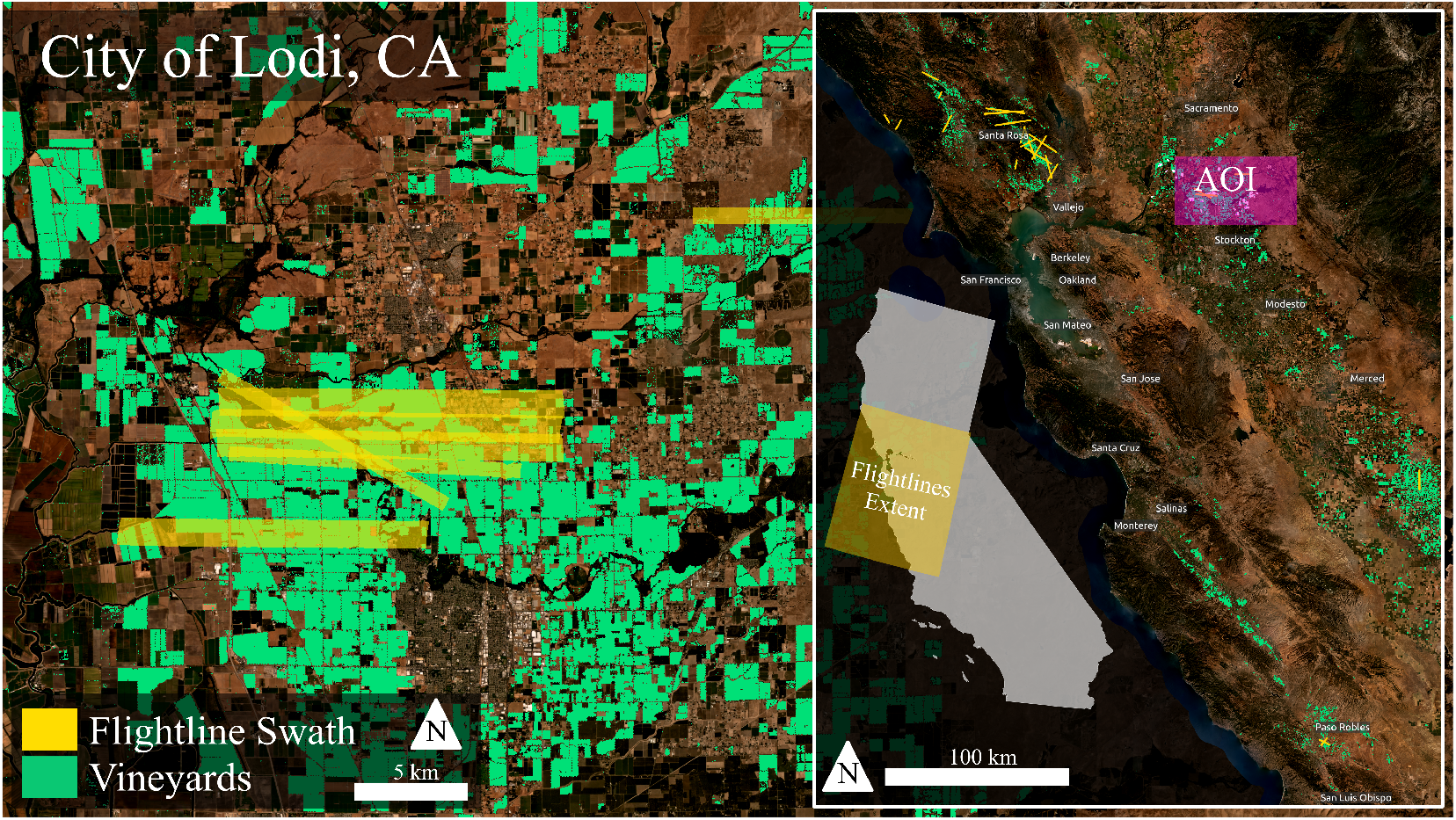
Region of interest (ROI) highlighting AVIRIS-NG acquisitions which spatially-overlap our ground validation data in the city of Lodi, California.

**Figure 3.**
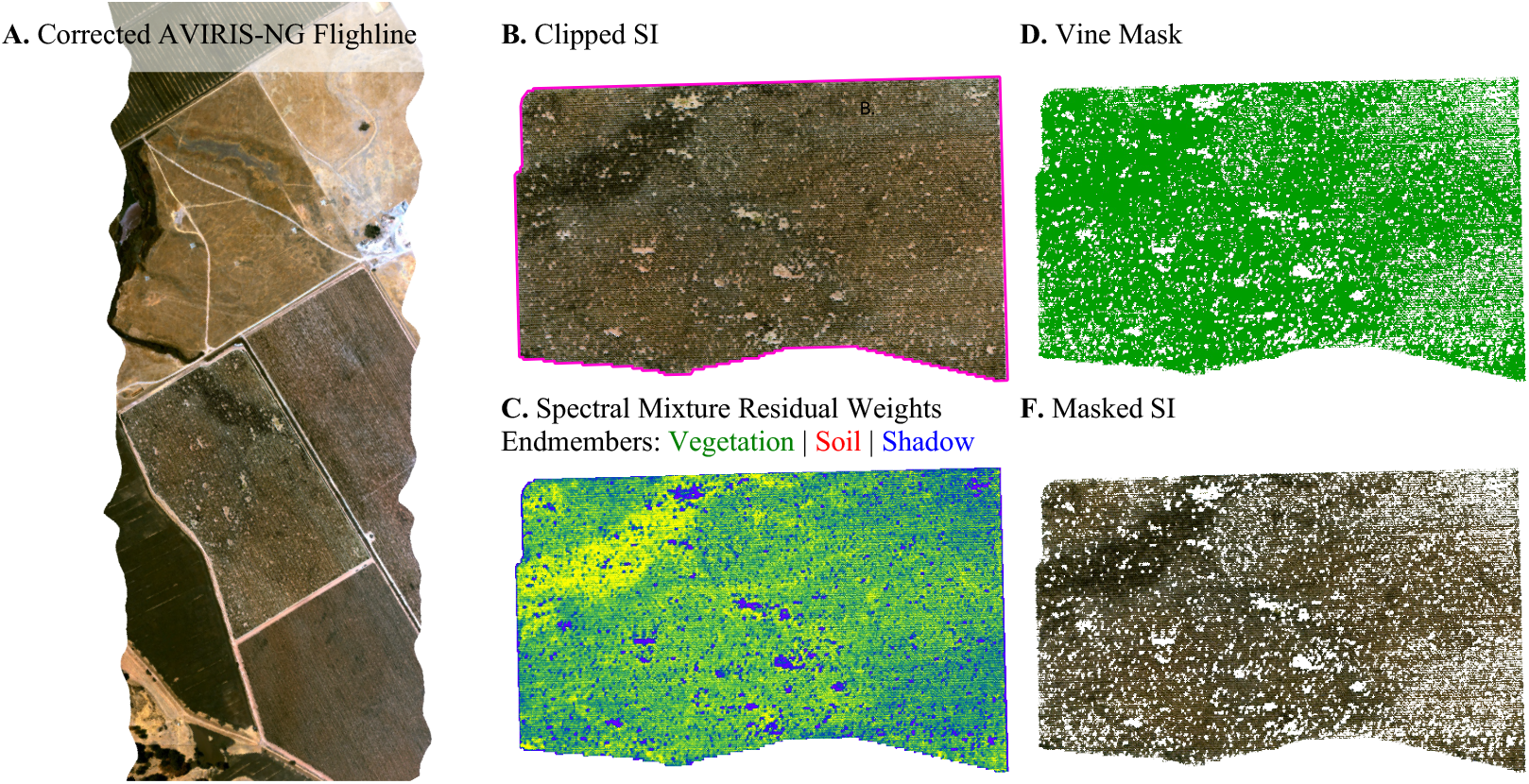
Step 0: **A.** Full AVIRIS-NG corrected SI. Step 1: **B.** SI clipped to the boundary of the vineyard. Step 2: **C.** SMR endmember weights. Step 3: **D.** Vine mask generated from the SMR weights. Step 4: **F.** Resulting SI image from pipeline.

## 3 Results

### 3.1 Experimental lab

The training at the experimental lab yields GLRaV-3 models that may be deployed locally or stored in the cloud for later retrieval and disease inference requests. In order to store the models in the cloud, they must be serialized (a process also known as ‘pickling’). Serialization is the process of turning objects in a computer program into bits that can be sent on a network like the Internet and/or stored in persistent storage like a hard disk drive (HDD) or solid-state drive (SSD). The models essentially represent mathematical functions that represent the most salient components in the disease detection process. Therefore, the serialization process captures these functions as (Python) objects that can be uploaded from one site and downloaded at another site with the same representation.

The model upload process (also known as registering the model in Figure 4) specifies the location of the model on the local (Grapekiller) computer, the name for the model, any appropriate tags, a short description, and the Azure Machine Learning (ML) workspace where the model is to be uploaded. Azure ML is a platform-as-a-service (PaaS) offering to enable big data processing in the Azure Cloud, and a workspace is a logical partition of computing and storage servers for a single project (Microsoft, 2021). Therefore, the name of the model must be unique within a workspace. The tags are useful to filter models during the download process. The short description is optional, but it is useful in case the model is to be shared across institutions. Within the workspace, models are indexed by names and versions. The versions are automatically created by the Azure ML workspace when an existing model name is registered more than once. In other words, all models are stored under a single location, but it is possible to specify which version to download and apply for inference requests.

**Figure 4.**
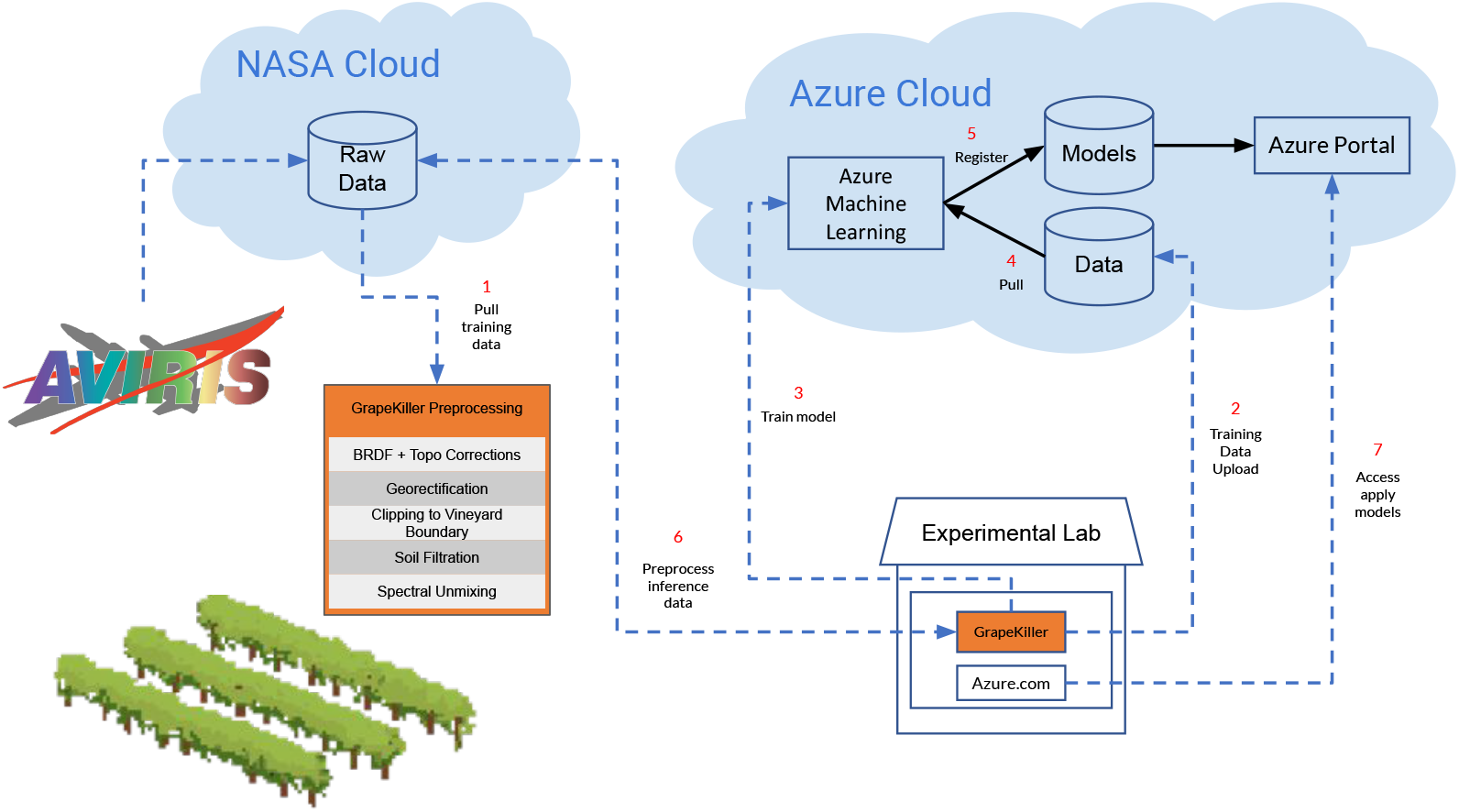
A software architecture for early disease detection

**Figure 5.**
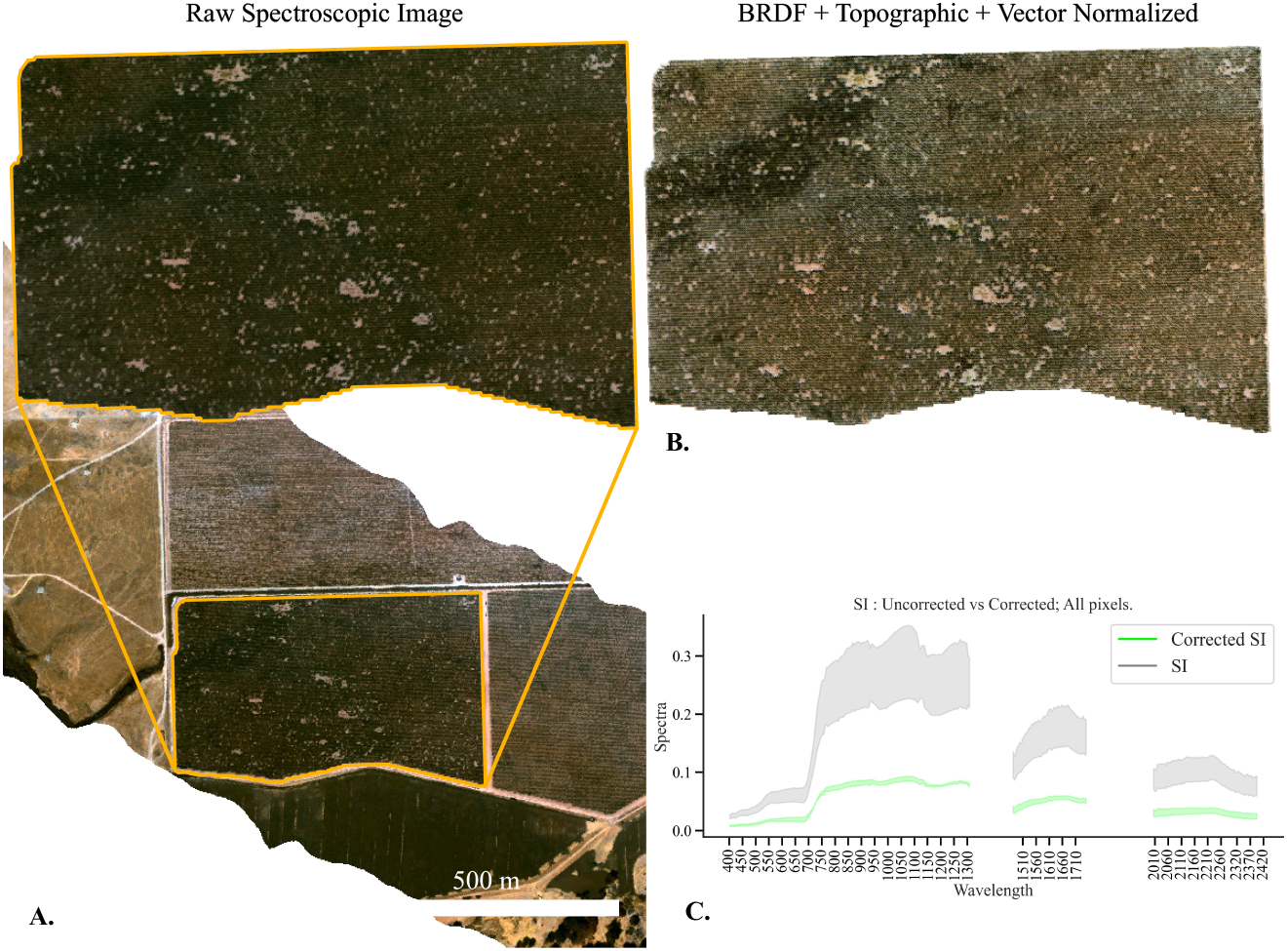
**A.** AVIRIS-NG Flightline with inset of example vineyard as seen by AVIRIS-NG. **B.** Imagery with BRDF, Topographic, and Vector normalization applied. **C.** Spectroscopic graph of reflectance values before and after.

The download process can be triggered programmatically or manually. The programmatic download can be triggered from any (Python) script with the correct credentials for the workspace where the model is stored. The programmatic trigger is ideal for downloading the model at the network edge where connectivity to the cloud is limited in order to field inference requests even during network outages. As demonstrated in step 7 of Figure 4, the manual download is possible from any networked computer with access credentials for the Azure ML workspace through the Azure portal. The Azure portal is an end-user-friendly interface for provisioning and managing resources in the Azure Cloud. The manual download is a convenient way to access the model parameters and potentially check the size before targeting a model deployment location or device. This is important for inference requests that may be run on resource-constrained devices such as Raspberry Pis (Raspberry Pi, 2022).

### 3.2 GLRaV-3 Detection Maps

The resulting classified rasters in Figure 6 are the final output from the model management pipeline. These rasters include the relevant disease incidence and stage within a three-meter pixel. The pipeline outputs six rasters: A classified raster for non-infected vine locations and infected vines (whether symptomatic or asymptomatic), a classified raster for non-infected vine locations and symptomatic vines, a classified raster for non-infected vine locations and asymptomatic vine locations, and the class probability for each respective raster.

**Figure 6.**
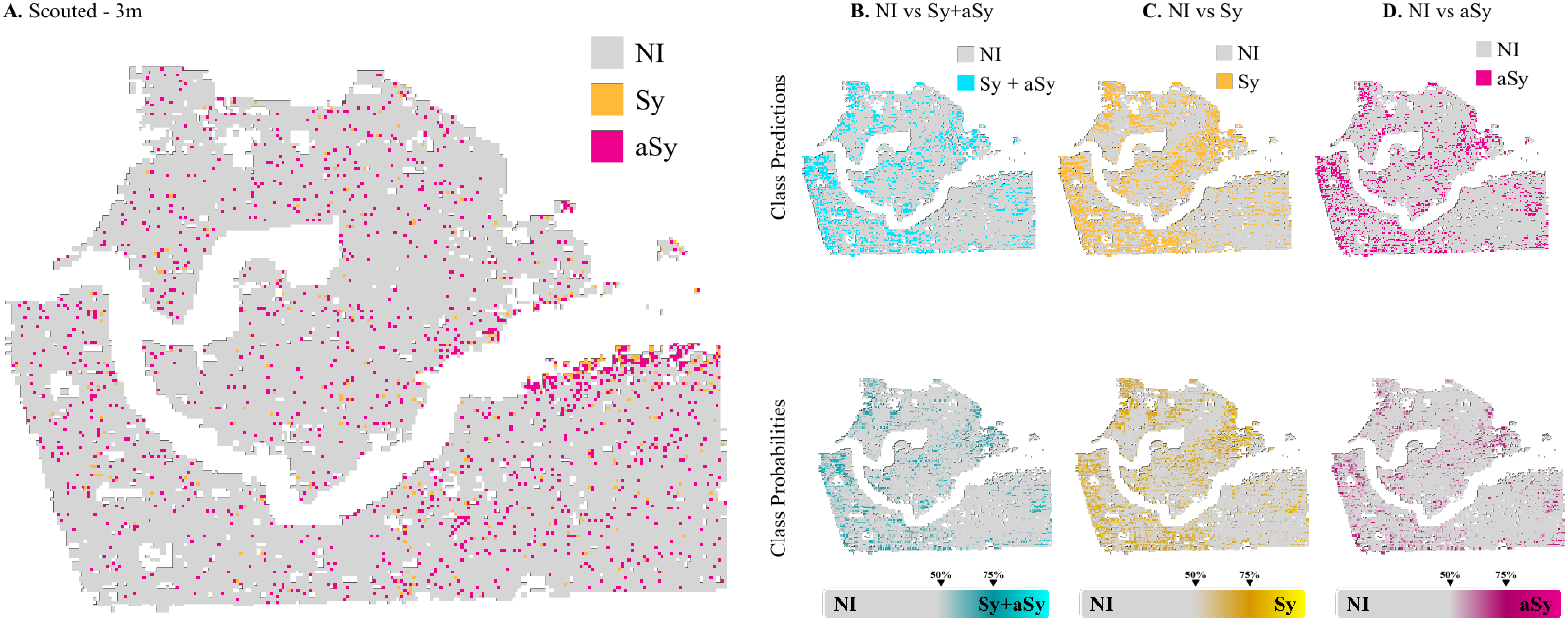
**A.** Scouted disease incidence. **B.** Predicted disease incidence of non-infected vs symptomatic or asymptomatic vines (top) and probability for each class (bottom). **C.** Predicted disease incidence raster for non-infected vs symptomatic vines (top) and probability for each class (bottom). **D.** Predicted disease incidence raster for non-infected vs asymptomatic vines (top) and probability for each class (bottom). Figure taken from (Romero Galvan et al., 2022)

## 4 Discussion

Our methodology and results showcase a robust system with significant implications for decision support not only for crop disease detection but also for future data modalities and use cases targeted to improve our understanding of biological Earth system processes. First, the proposed system architecture is general enough to enable plug-and-play of different and complementary models. For example, this architecture can easily be reused to train and test grape variety classification models, which in turn we can better refine our methodologies for disease detection across grape varieties. This adaptability may prove important for both future application and model improvement, given different management requirements and innate disease resistance across grape varieties which may affect the spectroscopic signal, as illustrated in Figure 7. Second, beyond model variety, we designed the software architecture to be plug-and-play to different data sources. Currently, the system provides GLRaV-3 decision support at 3m resolution with SI derived from AVIRIS-NG. As we look to future satellite missions, such as SBG and CHIME, our system can be leveraged to explore disease detection efficacy at these systems’ anticipate resolutions (30m) years ahead of their launches.

**Figure 7.**
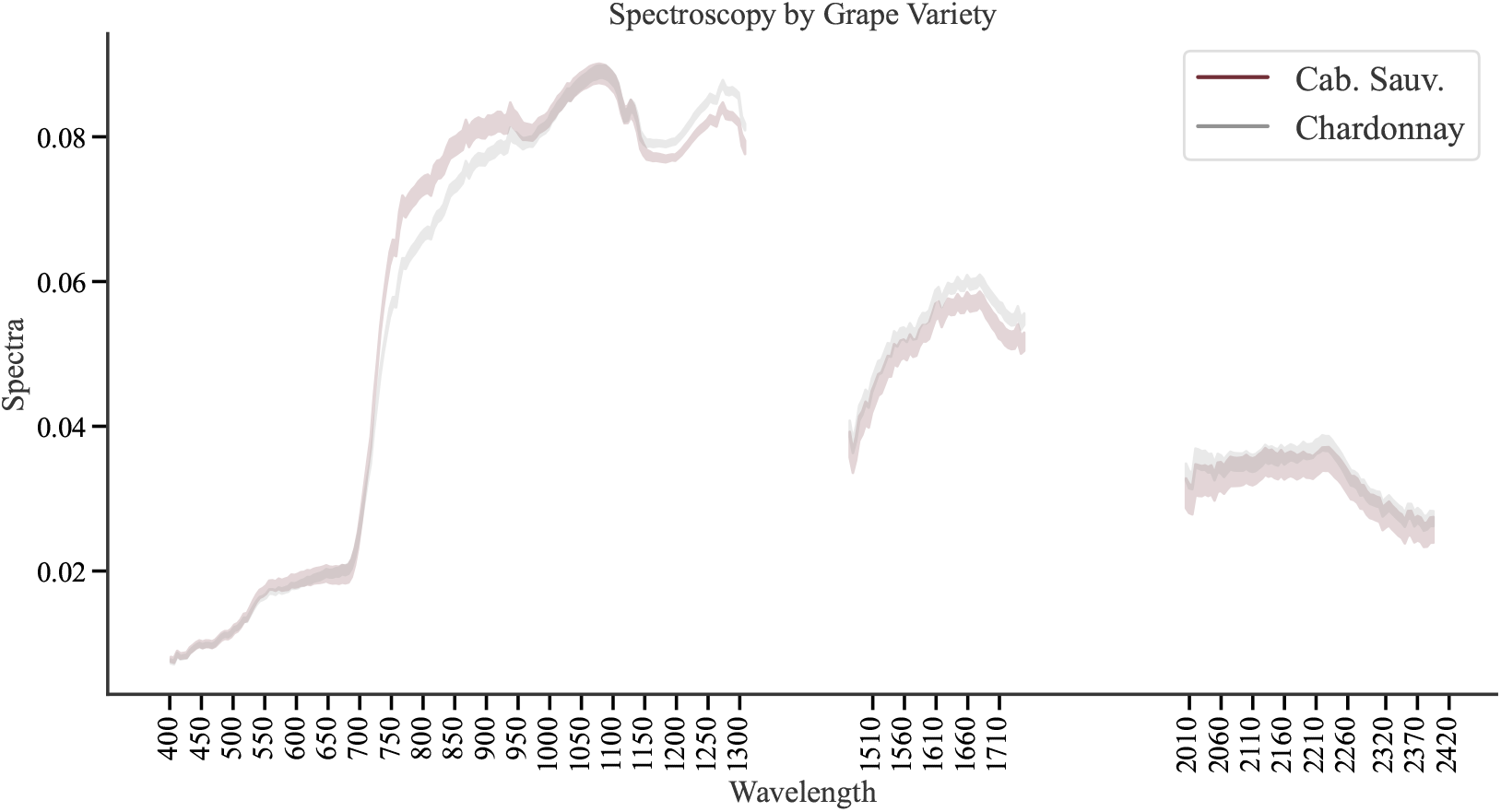
Spectroscopic signal of red-grape variety Cabernet Sauvignon vs white-grape variety Chardonnay.

In our context, the public cloud provides storage and computation spaces for the disease detection data and models (both of which are refined by the experimental lab) (Mell & Grance, 2011). It is important to note that although we use the Azure Cloud, the “Google Cloud”, “Amazon Cloud”, and “IBM Cloud”, could have been used as well. Our use of cloud computing plays a crucial role in fulfilling our architecture goals to have social impact and research reproducibility while preserving user privacy. Social impact is achieved by demonstrating a holistic, simple architecture that agricultural stakeholders can access as a service in the decision-making process via refined models. The models can be deployed as web endpoints that can be called upon to perform field data inference requests on edge devices. Our architecture also provides a platform that contributes to reproducible research while addressing user data privacy concerns by leaving all potentially proprietary data on the user’s edge device. Through the Azure ML platform that the architecture wraps around, researchers can access and share disease detection models within both their community of peers and stakeholders (Microsoft, 2021). As discussed in Section 3.1, Azure ML enables systematic registration, tracking, and tagging of models by researchers in the course of the experimentation process that is at the core of the scientific method. From iterative and tagged models, researchers can confidently make general and reproducible knowledge claims. While this architecture makes strides towards reproducibility, streamlined sharing of models (e.g. access permissions, appropriate application programming interfaces, or APIs) are still needed to enable true access and sharing not only across institutions but also across cloud providers. This challenge is not addressed in this work, and it remains an open area of exploration for our future work in this vein (see Section 4.1).

### 4.1 Limitations

Co-occurring biotic and abiotic stresses can confuse SI-informed GLRaV-3 detection (Romero Galvan et al., 2022). It is not uncommon for a vine, or multiple neighboring vines, to be infected with multiple pathogens (e.g. trunk disease), and/or experience co-occurring abiotic stress (e.g. drought). Further work is required to gauge how common abiotic stressors in Californian vineyards (e.g. water stress, insect damage, or sunburn) cause misclassifications. However, as long as the model correctly identifies a plant as abnormal, this information is still of great value to agricultural stakeholders because it can be used to strategically deploy scouts and field management teams to areas most in need of attention. Thus the potential service outlined here is still valuable to the grape grower community, despite this open area for improvement. Another limitation is AVIRIS-NG acquisition availability. While the majority of grapevine in the United States is in California, the home of AVIRIS-NG and thus the state with the most historical coverage, there are growers in other regions including New York, which is the 3rd most important grape-growing state in the US. AVIRIS-NG is an airborne platform that requires appropriate flying conditions for an acquisition to happen, and thus can be both expensive and logistically challenging to deploy. However, spaceborne imaging spectrometers such as SBG, Soil Mapper, and CHIME will soon provide global SI. With the help of this proposed framework, we open the door for global-scale GLRaV-3 monitoring beyond the currently outlined geographic area of study.

In this work, we have demonstrated the efficacy of our Azure Cloud-based disease detection system. The next logical step is to streamline the system toward a more accessible user experience for agricultural stakeholders and researchers. However, we face two limitations. First, although Azure provides the capability to deploy the models as web-based endpoints (e.g. issuing web requests with shape files and getting back classified rasters of disease presence), the disease detection system is still tied to a single cloud provider. To avoid technical compatibility issues inherent in vendor lock-in (e.g. transferring data across private and public clouds), the system must evolve to work across clouds so that the data and models are not siloed in a single cloud. Second, the system’s effectiveness has been shown in a few fields. In practice, growers operate thousands of fields potentially spread across multiple farms. One area that remains unexplored is managing diverse models across farms. For instance, growers can operate water-stress models at some farms while deploying GLRaV-3 models on other fields. Technologically, this necessitates scalable techniques for managing the model and data lifecycle across farms and is the subject of ongoing future work.

## 5 Conclusions

Here, we describe and deploy an adaptable cloud-based system for detecting an economically important crop disease, GLRaV-3 in grapevine, with airborne imaging spectroscopy data. The proposed architecture only requires SI from an airborne or spaceborne source and a shapefile from the user specifying location coordinates and boundaries to run. The goal of this system is to empower agricultural stakeholders to make well-informed, data-driven decisions by granting them access to the latest advances in SI-ML disease detection. Our work improves crop disease decision-making while serving as a guide for others interested in developing accessible, use-inspired, remote sensing applications for agricultural stakeholders who stand to benefit from publicly funded research.

Besides the exploration of different models and data modalities, the system could fundamentally change the way we approach vineyard management operations. Infected vines, be they symptomatic or asymptomatic, still produce grapes, albeit of lower quality for the wine-making process, which chips away at profit across the value chain. Typically, asymptomatic vines are hard to detect, which ultimately affects grape and wine quality. The system presented in this work can provide a valuable predictor/indicator of potential yield, as well as which vines are spectroscopic anomalies, signaling stress, disease, or any other damage. This predictive analytics service is useful in anticipating the chemical supplementation that may be necessary for the upcoming harvest and fermentation of lesser-quality grapes. From a financial and labor perspective, molecular testing, the most reliable method of asymptomatic detection, does not scale to the sizes required. However, armed with a disease detection system, growers can strategically deploy their high-accuracy ground resources, such as expert scouts or molecular testing, more efficiently because the system predicts with high accuracy vineyard regions most likely to be infected. In this manner, the system complements the farm ecosystem, empowering the users, and not seeking to replace their valuable expertise.

## 6 Open Research

All spectroscopic imagery used for this study are publicly available without restrictions from the AVIRIS-NG Data portal: https://aviris.jpl.nasa.gov/dataportal/. Random forest models developed for detecting GLRaV-3 are hosted on a public GitHub repository, available here: https://github.com/GoldLab-GrapeSpec/GLRaV3Detection. Likewise, python scripts are used to apply the models and anaconda environment to execute the scripts.

## 7 Acknowledgements

This work was funded in part by NASA FINESST (Grant 80NSSC21K1605) awarded to Romero Galvan (FI) and Gold (PI), NASA Jet Propulsion Laboratory Strategic University Research Partnership Fund, NSF NRT Digital Plant Sciences training grant awarded to Cornell University (Grant 1922551), the NSF STC Center for Research on Programmable Plant Systems (CROPPS; Grant 2019674), and the NSF CHS grant understanding and improving the social impact of high-bandwidth farm networking infrastructure (Grant number 1955125). We acknowledge with gratitude Dr. Michael Scanlon, project leader of the Cornell University NSF NRT Digital Plant Sciences program that introduced and inspired Rubambiza and Romero Galvan to collaborate on this co-first authored paper. Additionally, we thank Dr. Michael Eastwood and the myriad members of the NASA JPL AVIRIS-NG team who, along with the NASA Biodiversity and Ecological Conservation program office, are responsible for initiating and executing the successful air campaign that provided data for this publication. Importantly, we would like to thank our industry collaborators, including Charlie Starr, Stephanie Bolton, Mimar Alsina, and Nick Dokoozlian, for their invaluable efforts, feedback, and collaboration that supported this project.

## References

Almeida, R. P. P., Daane, K. M., Bell, V. A., Blaisdell, G. K., Cooper, M. L., Herrbach, E., & Pietersen, G. (2013). Ecology and management of grapevine leafroll disease. Frontiers in Microbiology, 4. Retrieved 2022-10-02, from http://journal.frontiersin.org/article/10.3389/fmicb.2013.00094/abstract doi: 10.3389/fmicb.2013.00094

Amazon Web Services, I. (2022, November). NFL Next Gen Stats Powered by AWS - Stat That. Retrieved 2022-11-17, from https://aws.amazon.com/sports/nfl/

Analytics, A. (2022, November). Ag-Analytics: Discover Agricultural Data. Retrieved 2022-11-13, from https://www.analytics.ag

Chapman, J. W., Thompson, D. R., Helmlinger, M. C., Bue, B. D., Green, R. O., Eastwood, M. L., … Lundeen, S. R. (2019, September). Spectral and Radiometric Calibration of the Next Generation Airborne Visible Infrared Spectrometer (AVIRIS-NG). Remote Sensing, 11 (18), 2129. Retrieved 2022-09-06, from https://www.mdpi.com/2072-4292/11/18/2129 doi: 10.3390/rs11182129

Charles, J., Cohen, D., Walker, J., Forgie, S., Bell, V., & Breen, K. (2006, August). A review of the ecology of grapevine leafroll associated virus type 3 (GLRaV3). New Zealand Plant Protection, 59, 330–337. Retrieved 2022-09-06, from https://nzpps.org/_journal/index.php/nzpp/article/view/4590 doi: 10.30843/nzpp.2006.59.4590

Charles, J. G., Froud, K. J., van den Brink, R., & Allan, D. J. (2009). Mealybugs and the spread of grapevine leafroll-associated virus 3 (GLRaV-3) in a New Zealand vineyard. Australasian Plant Pathology, 38(6), 576. Retrieved 2022-09-06, from http://link.springer.com/10.1071/AP09042 doi: 10.1071/AP09042

Chlus, A., Winstonolson, Greenberg, E., & Oldmanye007. (2022, August). EnSpec/hytools: 1.4.0. Zenodo. Retrieved 2022-09-06, from https://zenodo.org/record/5997755 (Language: en) doi: 10.5281/ZENODO.5997755

Cohen, J. (2021, December). 4 Companies Control 67% of the World’s Cloud Infrastructure. Retrieved 2022-11-13, from https://www.pcmag.com/news/four-companies-control-67-of-the-worlds-cloud-infrastructure

Curran, P. J. (1989, December). Remote sensing of foliar chemistry. Remote Sensing of Environment, 30(3), 271–278. Retrieved 2022-09-06, from https://linkinghub.elsevier.com/retrieve/pii/0034425789900692 doi: 10.1016/0034-4257(89)90069-2

Gao, B.-C., & Goetz, A. F. H. (1990). Column atmospheric water vapor and vegetation liquid water retrievals from Airborne Imaging Spectrometer data. Journal of Geophysical Research, 95(D4), 3549. Retrieved 2022-11-16, from http://doi.wiley.com/10.1029/JD095iD04p03549 doi: 10.1029/JD095iD04p03549

Gholamy, A., Kreinovich, V., & Kosheleva, O. (2018). Why 70/30 or 80/20 relation between training and testing sets: A pedagogical explanation..

Gillies, S., et al. (2013–). Rasterio: geospatial raster i/o for Python programmers. Retrieved from https://github.com/rasterio/rasterio

Golino, D. A., Sim, S. T., Gill, R., & Rowhani, A. (2002, November). California mealybugs can spread grapevine leafroll disease. California Agriculture, 56(6), 196–201. Retrieved 2022-09-06, from https://calag.ucanr.edu/archive/?article=ca.v056n06p196 doi: 10.3733/ca.v056n06p196

Google, I. (2022, November). Cloud Computing Services. Retrieved 2022-11-18, from https://cloud.google.com/

Hruŝka, J., Adão, T., Pádua, L., Marques, P., Peres, E., Sousa, A., … Sousa, J. J. (2018). Deep learning-based methodological approach for vineyard early disease detection using hyperspectral data. In Igarss 2018 - 2018 ieee international geoscience and remote sensing symposium (p. 9063–9066). doi: 10.1109/IGARSS.2018.8519136

IBM. (2022, October). IBM Cloud. Retrieved 2022-11-18, from https://www.ibm.com/cloud

Jiménez-Brenes, F. M., López-Granados, F., Torres-Sánchez, J., Peña, J. M., Ramírez, P., Castillejo-González, I. L., & de Castro, A. I. (2019). Automatic uav-based detection of cynodon dactylon for site-specific vineyard management. PloS one, 14 (6), e0218132.

Maree, H. J., Almeida, R. P. P., Bester, R., Chooi, K. M., Cohen, D., Dolja, V. V., … Burger, J. T. (2013). Grapevine leafroll-associated virus 3. Frontiers in Microbiology, 4. Retrieved 2022-10-02, from http://journal.frontiersin.org/article/10.3389/fmicb.2013.00082/abstract doi: 10.3389/fmicb.2013.00082

Marr, B. (2015, September). Big Data: 20 Mind-Boggling Facts Everyone Must Read. Retrieved 2022-11-15, from https://www.forbes.com/sites/bernardmarr/2015/09/30/big-data-20-mind-boggling-facts-everyone-must-read/

Mell, P., & Grance, T. (2011). The NIST Definition of Cloud Computing (Tech. Rep.). Gaithersburg, MD, USA: National Institute of Standards and Technology.

Microsoft. (2021, May). What is an Azure Machine Learning workspace. Retrieved n.d., from https://www.twilio.com/docs/sms/quickstart/pytho://docs.microsoft.com/en-us/azure/machine-learning/concept-workspace

Microsoft. (2022, November). What is Azure—Microsoft Cloud Services — Microsoft Azure. Retrieved 2022-11-18, from https://azure.microsoft.com/en-us/resources/cloud-computing-dictionary/what-is-azure/

Naidu, R., Rowhani, A., Fuchs, M., Golino, D., & Martelli, G. P. (2014, September). Grapevine Leafroll: A Complex Viral Disease Affecting a High-Value Fruit Crop. Plant Disease, 98(9), 1172–1185. Retrieved 2022-09-06, from https://apsjournals.apsnet.org/doi/10.1094/PDIS-08-13-0880-FE doi: 10.1094/PDIS-08-13-0880-FE

Nieke, J., & Rast, M. (2018, July). Towards the Copernicus Hyperspectral Imaging Mission For The Environment (CHIME). In IGARSS 2018 - 2018 IEEE International Geoscience and Remote Sensing Symposium (pp. 157–159). Valencia: IEEE. Retrieved 2022-11-11, from https://ieeexplore.ieee.org/document/8518384/ doi: 10.1109/IGARSS.2018.8518384

Nnadi, N. E., & Carter, D. A. (2021, April). Climate change and the emergence of fungal pathogens. PLOS Pathogens, 17(4), e1009503. Retrieved 2022-11-11, from https://dx.plos.org/10.1371/journal.ppat.1009503 doi: 10.1371/journal.ppat.1009503

Olmos, A., Bertolini, E., Ruiz-García, A. B., Martínez, C., Peiró, R., & Vidal, E. (2016, May). Modeling the Accuracy of Three Detection Methods of *Grapevine leafroll-associated virus 3* During the Dormant Period Using a Bayesian Approach. Phytopathology®, 106(5), 510–518. Retrieved 2022-09-06, from https://apsjournals.apsnet.org/doi/10.1094/PHYT0-10-15-0246-R doi: 10.1094/PHYTO-10-15-0246-R

Ortiz-Bobea, A., Ault, T. R., Carrillo, C. M., Chambers, R. G., & Lobell, D. B. (2021, April). Anthropogenic climate change has slowed global agricultural productivity growth. Nature Climate Change, 11 (4), 306–312. Retrieved 2022-11-11, from http://www.nature.com/articles/s41558-021-01000-1 doi: 10.1038/s41558-021-01000-1

Pietersen, G., Spreeth, N., Oosthuizen, T., van Rensburg, A., van Rensburg, M., Lottering, D., … Tooth, D. (2013, June). Control of Grapevine Leafroll Disease Spread at a Commercial Wine Estate in South Africa: A Case Study. American Journal of Enology and Viticulture, 64 (2), 296–305. Retrieved 2022-09-06, from http://www.ajevonline.org/cgi/doi/10.5344/ajev.2013.12089 doi: 10.5344/ajev.2013.12089

Queally, N., Ye, Z., Zheng, T., Chlus, A., Schneider, F., Pavlick, R. P., & Townsend, P. A. (2022, January). FlexBRDF: A Flexible BRDF Correction for Grouped Processing of Airborne Imaging Spectroscopy Flightlines. Journal of Geophysical Research: Biogeosciences, 127(1). Retrieved 2022-09-06, from https://onlinelibrary.wiley.com/doi/10.1029/2021JG006622 doi: 10.1029/2021JG006622

Raspberry Pi, L. (2022, November). Buy a Raspberry Pi 4 Model B. Retrieved 2022-11-16, from https://www.raspberrypi.com/products/raspberry-pi-4-model-b/

Romero Galvan, F. E., Pavlick, R., Trolley, G., Aggarwal, S., Sousa, D., Starr, C., … Gold, K. M. (2022, October). Scalable early detection of grapevine virus infection with airborne imaging spectroscopy (preprint). Pathology. Retrieved 2022-11-14, from http://biorxiv.org/lookup/doi/10.1101/2022.10.04.510827 doi: 10.1101/2022.10.04.510827

Sapes, G., Lapadat, C., Schweiger, A. K., Juzwik, J., Montgomery, R., Gholizadeh, H., … Cavender-Bares, J. (2022, May). Canopy spectral reflectance detects oak wilt at the landscape scale using phylogenetic discrimination. Remote Sensing of Environment, 273, 112961. Retrieved 2022-09-06, from https://linkinghub.elsevier.com/retrieve/pii/S003442572200075X doi: 10.1016/j.rse.2022.112961

Scheffler, D., Hollstein, A., Diedrich, H., Segl, K., & Hostert, P. (2017, July). AROSICS: An Automated and Robust Open-Source Image Co-Registration Software for Multi-Sensor Satellite Data. Remote Sensing, 9(7), 676. Retrieved 2022-09-06, from http://www.mdpi.com/2072-4292/9/7/676 doi: 10.3390/rs9070676

Schneider, F., Ferraz, A., & Schimel, D. (2019, October). Watching Earth’s Interconnected Systems at Work. Eos, 100. Retrieved 2022-11-14, from https://eos.org/science-updates/watching-earths-interconnected-systems-at-work doi: 10.1029/2019EO136205

Sousa, D., Brodrick, P., Cawse-Nicholson, K., Fisher, J. B., Pavlick, R., Small, C., & Thompson, D. R. (2022, February). The Spectral Mixture Residual: A Source of Low-Variance Information to Enhance the Explainability and Accuracy of Surface Biology and Geology Retrievals. Journal of Geophysical Research: Biogeosciences, 127(2). Retrieved 2022-09-06, from https://onlinelibrary.wiley.com/doi/10.1029/2021JG006672 doi: 10.1029/2021JG006672

Thompson, D. R., Natraj, V., Green, R. O., Helmlinger, M. C., Gao, B.-C., & Eastwood, M. L. (2018, October). Optimal estimation for imaging spectrometer atmospheric correction. Remote Sensing of Environment, 216, 355–373. Retrieved 2022-10-02, from https://linkinghub.elsevier.com/retrieve/pii/S0034425718303304 doi: 10.1016/j.rse.2018.07.003

Zarco-Tejada, P. J., Camino, C., Beck, P. S. A., Calderon, R., Hornero, A., Hernández-Clemente, R., … Navas-Cortes, J. A. (2018, July). Previsual symptoms of Xylella fastidiosa infection revealed in spectral plant-trait alterations. Nature Plants, 4 (7), 432–439. Retrieved 2022-09-06, from http://www.nature.com/articles/s41477-018-0189-7 doi: 10.1038/s41477-018-0189-7

Zarco-Tejada, P. J., Poblete, T., Camino, C., Gonzalez-Dugo, V., Calderon, R., Hornero, A., … Navas-Cortes, J. A. (2021, December). Divergent abiotic spectral pathways unravel pathogen stress signals across species. Nature Communications, 12(1), 6088. Retrieved 2022-09-06, from https://www.nature.com/articles/s41467-021-26335-3 doi: 10.1038/s41467-021-26335-3

